# Cholecystokinin Suppresses β-Cell Apoptosis, Including in Human Islets in a Transplant Model

**DOI:** 10.1101/2021.03.02.433645

**Authors:** Hung Tae Kim, Arnaldo H. de Souza, Heidi Umhoefer, JeeYoung Han, Lucille Anzia, Steven J. Sacotte, Rashaun A. Williams, Joseph T. Blumer, Jacob T. Bartosiak, Danielle A. Fontaine, Mieke Baan, Carly R. Kibbe, Dawn Belt Davis

**Author notes:** Co-First Authors (Hung Tae Kim, Arnaldo H. de Souza). Co-Corresponding Authors: Dawn. B. Davis –, Carly. R. Kibbe –.

## Abstract

Loss of functional pancreatic β-cell mass and increased β-cell apoptosis are fundamental to the pathophysiology of both type 1 and type 2 diabetes. Pancreatic islet transplantation has the potential to cure type 1 diabetes but is often ineffective due to the death of the islet graft within the first few years after transplant. Therapeutic strategies to directly target pancreatic β-cell survival are needed to prevent and treat diabetes and to improve islet transplant outcomes. Reducing β-cell apoptosis is also a therapeutic strategy for type 2 diabetes. Cholecystokinin (CCK) is a peptide hormone typically produced in the gut after food intake, with positive effects on obesity and glucose metabolism in mouse models and human subjects. We have previously shown that pancreatic islets also produce CCK. The production of CCK within the islet promotes β-cell survival in rodent models of diabetes and aging. Now, we demonstrate a direct effect of CCK to reduce cytokine-mediated apoptosis in a β-cell line and in isolated mouse islets in a receptor-dependent manner. However, whether CCK can protect human β-cells was previously unknown. Here, we report that CCK can also reduce cytokine-mediated apoptosis in isolated human islets and CCK treatment *in vivo* decreases β-cell apoptosis in human islets transplanted into the kidney capsule of diabetic NOD/SCID mice. Collectively, these data identify CCK as a novel therapy that can directly promote β-cell survival in human islets and has therapeutic potential to preserve β-cell mass in diabetes and as an adjunct therapy after transplant.

**One Sentence Summary:** Cholecystokinin ameliorates pancreatic β-cell death under models of stress and after transplant of human islets.

## INTRODUCTION

Diabetes mellitus (DM) is a multifactorial disease that results from impairment of β-cell function and loss of β-cell mass. Type 1 DM (T1D) develops from autoimmune destruction of the pancreatic β-cells, resulting in insulin dependence ^1^. Type 2 DM (T2D) results from both increased insulin resistance and decreased insulin production ^2^. There is a failure of adaptive pancreatic β-cell proliferation under diabetic stress, as well as increased apoptosis leading to β-cell mass reduction^2,3^. Epidemiological studies show that during the past three decades, there has been an alarming rate of growth in the prevalence of both types of diabetes, consequently driving diabetes-related complications and mortality ^4^. In 2020, the American Diabetes Association reported that 34.2 million Americans are diagnosed with diabetes, while 88 million more Americans have prediabetes ^5^. Yet, there are no treatment strategies specifically designed to protect against a decrease in pancreatic islet mass and prevent the loss of insulin producing cells ^6–8^. Many patients with T2D eventually require insulin therapy due to a decline in functional β-cell mass, yet exogenous insulin delivery cannot replicate the dynamic insulin production of a functional islet and dysglycemia often persists ^9^.

One possible curative therapy for diabetes is transplantation of purified human pancreatic islets. The first human islet cell transplantation (ICT) took place in Minneapolis in 1974 ^10^ and with the landmark clinical trial of the Edmonton protocol in 1999, allogenic-ICT became a potential solution to restoring glucose homeostasis for type 1 diabetes (T1D), while auto-ICT became an option of preserving endocrine function for patients requiring total pancreatectomy for chronic pancreatitis ^11^. Considerable improvements in transplant methods were made over the past three decades. However, the major problem with any ICT, particularly for a hepatic intraportal graft which is the primary site for transplantation in humans, is the significant loss of islet viability both early and late after transplant ^12^. Allogenic ICT only results in about 25-50% insulin independence after five years. New therapeutic approaches to protect and preserve β-cell mass in transplanted islets are necessary to improve outcomes.

Cholecystokinin (CCK) is a peptide hormone produced in the small intestine and brain, stimulating the release of digestive enzymes from the exocrine pancreas and inducing satiety ^13^. A large body of literature from different fields including neuroscience, cardiology, immunology, and endocrinology documents that CCK has anti-inflammatory ^14–16^, proliferative ^17–19^, and anti-apoptotic ^20,21^ effects against various insults in a wide range of pathophysiologic models. Combined, there is accumulating evidence that CCK receptor signaling can attenuate the onset and progression of several modes of cell death in various mammalian cell types and different organs. Systemic injection of CCK receptor agonists were once extensively studied as a possible anti-obesity agent due to its potential modulation of leptin signaling ^22–24^ and its role in central nervous system-mediated appetite suppression ^25,26^. In the context of diabetes, systemic injection of CCK analog in humans, and rodents ^27–29^ has been shown to improve glucose tolerance. While this has previously been thought to be due to an ability for CCK to stimulate insulin secretion, we have recently shown that CCK does not directly enhance glucose-stimulated insulin secretion in mouse or human islets^30^. In addition to its role as a circulating endocrine hormone from the neuroendocrine cells of the intestinal epithelium, CCK production is highly upregulated in mouse pancreatic β-cells under conditions of obesity and insulin resistance and is also produced and secreted from human islets ^31–35^. Mice with β-cell-specific CCK overexpression are protected against β-cell death after injection of the β-cell toxin streptozotocin and have preserved β-cell mass with aging ^32^. Conversely, CCK null obese mice have decreased islet mass and increased β-cell apoptosis ^31^. In mice and murine Min6 cells ^36^, as well as in 1.1B4 human β-cell lines ^37^, exogenous cholecystokinin treatment had been demonstrated to have anti-apoptotic effects.

Despite the accumulated observations of a pro-survival effect of CCK in various cell types and its overall potential benefits in weight regulation and glucose homeostasis^46^ the ability of CCK to directly protect human pancreatic β-cells has not been previously tested. Here we show that CCK treatment results in protection against β-cell apoptosis in various models ranging from *in vitro* insulinoma cell lines and *ex vivo* intact mouse and human islet. We demonstrate that this is a direct effect of CCK on its receptors in the islet. Additionally, we demonstrate that CCK-8 can improve human islet graft survival following transplant. The translational impact of this study lies in the identification of CCK-based therapeutics as a viable target for preventing β-cell apoptosis in humans. This has relevance for T1D and T2D, including in the setting of transplant or β-cell replacement therapy.

## RESULTS

### CCK Treatment Protects INS1E Cells Against Cytokine-Induced Apoptosis

We have previously shown that the localized production of CCK in pancreatic β-cells leads to improved β-cell survival in mouse models of obesity, aging, and diabetes ^31–34^. However, in these experiments, locally produced intra-islet CCK could have indirect effects on other cell types to mediate its pro-survival effects. To look more specifically at the direct pro-survival effects of CCK peptide treatment on the β-cell *in vitro*, we first turned to the rat insulinoma cell line, INS1E. In all experiments described, we used a stable analogue of sulfated CCK-8, s-Glu-Gln-CCK-8, that has high biologic activity *in vitro* and *in vivo* ^27,28,38^. We will refer to this peptide simply as CCK or CCK-8 throughout the manuscript. A time course of INS1E cell viability over 72 hours of exposure to mouse pro-inflammatory cytokine cocktail (10 ng/ul IL-1β, 50 ng/ul IFN-γ, and 50 ng/ul TNFα) demonstrates that CCK-8 (100 nM) treatment mitigates cytokine-induced cell death, as measured by trypan blue exclusion, (n=7, *p* <0.05) up to 48 hours (Fig.1A). Viability of INS1E cells treated with CCK-8 was superior to those treated with GLP-1 (100nM), another peptide hormone with known pro-survival effects in β-cells *in vitro* and a widely used therapeutic target for type 2 diabetes (Fig.1A). We further demonstrate that CCK-8 also protected INS1E cells from cytokine-induced apoptosis in a concentration-dependent manner, using image flow cytometry with annexin V and propidium iodide staining to identify cells in various stages of apoptosis (Fig.1B, n=3-6, *p* <0.001).

**Fig 1.**
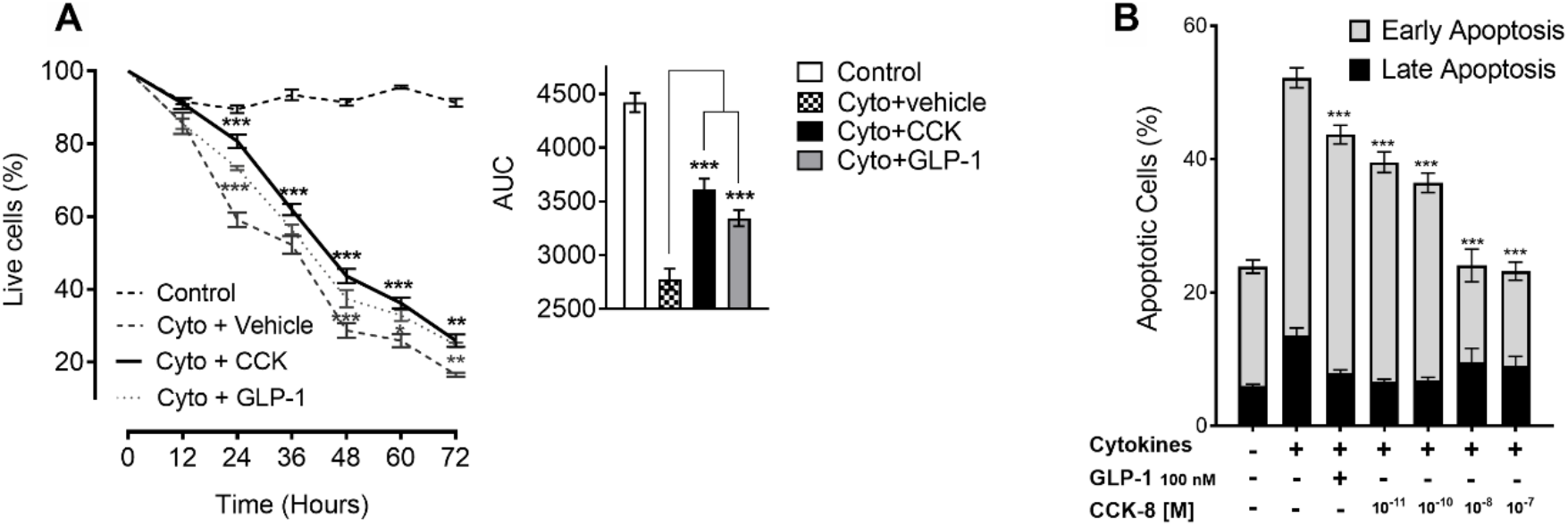
CCK protects INS1E cells from cytokine-induced apoptosis. Time-course (A) of INS1E cells treated with cytokine cocktail and co-treated with 100 nM CCK or 100 nM GLP-1 (n = 6-19). CCK dose-response effects (B) on the percentage of annexin V and propidium iodide (PI) double-positive cells (Late apoptosis – black bars) and Annexin V positive cells (Early Apoptosis - gray bars) treated with cytokines (n = 3-6). Data are means ± SEM; ***P < 0.001.

### CCK Treatment Protects Mouse Pancreatic β-Cells from Apoptosis

To determine whether CCK also has a direct effect on survival in primary mouse β-cells, we measured β-cell apoptosis of primary mouse islet cells by TUNEL assay. Intact mouse islets were pre-incubated with CCK-8 or vehicle for 24 hours. Mouse islets were then co-incubated with CCK-8 or vehicle and mouse cytokine cocktail (10 ng/ul IL-1β, 50 ng/ul IFN-γ, and 50 ng/ul TNFα) for an additional 24 hours. Mouse islets treated with CCK-8 (100 nM) had a 27% reduction in β-cell apoptosis after cytokine treatment compared to vehicle control, as measured by TUNEL staining (3.18% ± 0.27 CCK vs. 4.38% ± 0.29 vehicle vs. 0.36% ± 0.15 non-treated) (Fig.2A). In addition to the ability to protect from cytokine-mediated apoptosis, which is due to a combination of cell stressors, we wanted to test whether CCK also reduced β-cell apoptosis specifically due to endoplasmic reticulum (ER) stress. ER stress is elevated in β-cells in models of both type 1 and type 2 diabetes ^39,40^. Mouse islets that were treated with CCK-8 (100 nM, including pre-treatment for 24 hours) and thapsigargin (10 uM) to induce ER stress have significantly fewer TUNEL positive cells than cells treated with thapsigargin alone (Fig.2B). Finally, to demonstrate that the reduction in apoptosis was a direct, CCK-receptor mediated effect, we tested whether CCK-8 could still protect from apoptosis in islets from mice with knockout of both the CCKA and CCKB receptors (ABKO). We demonstrate using propidium iodide and annexin V staining via imaging flow cytometry that the protective effect of CCK-8 (100 nM) seen in the WT islets disappears in islets from CCKABR KOs, indicating that protection of mouse islets against cytokines by CCK is CCK receptor-dependent (Fig.2C). Taken together, these findings suggest that activation of CCK receptors protects rodent β-cells from apoptosis due to multiple stressors directly through the CCKAR, CCKBR, or a combination of both receptors.

**Fig 2.**
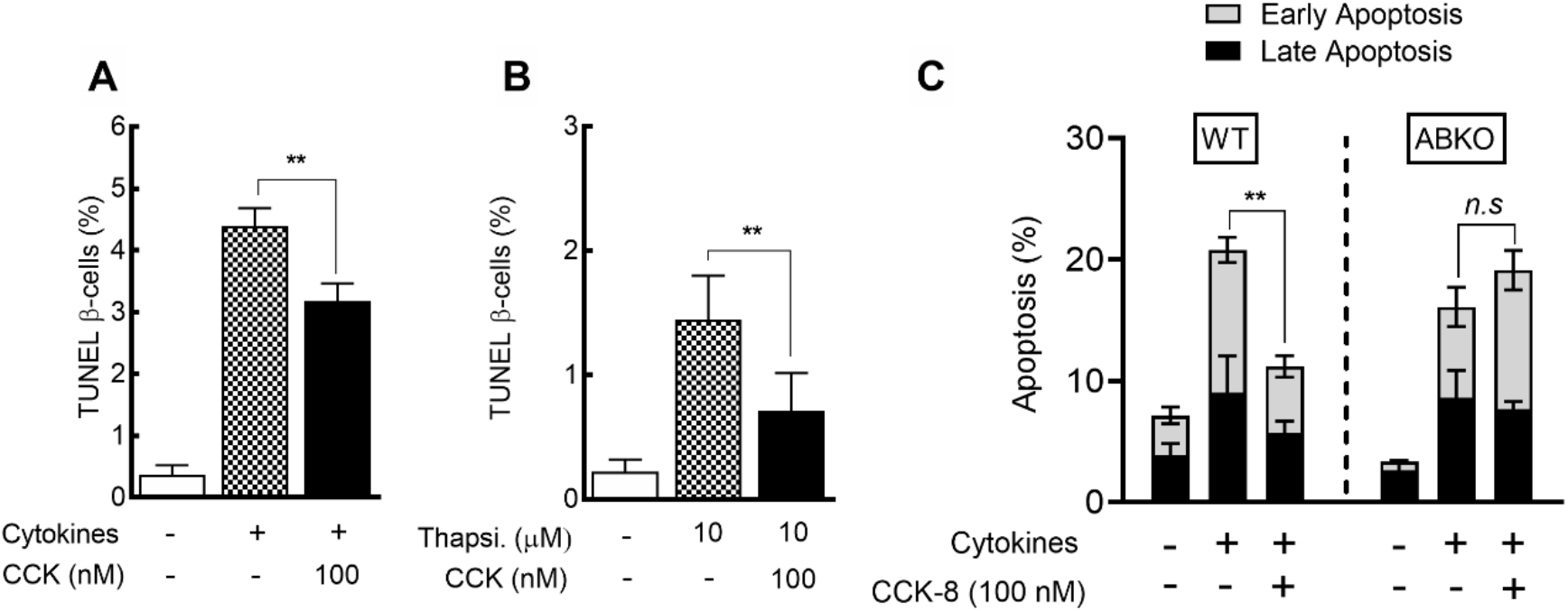
CCK protects mouse islets from apoptosis. Percentage of TUNEL-positive insulin-positive cells of mouse islets treated with cytokine cocktail (A) or thapsigargin (B) is reduced when treated with 100 nM CCK (*n* = 3-4). (C) Percentage of Annexin V and propidium iodide (PI) double-positive cells (Late apoptosis - black bars) and Annexin V positive mouse islet cells (Early Apoptosis - gray bars) co-treated with cytokines and CCK is reduced in wild type controls but not in double CCK receptor KO islets (ABKO) (*n* = 6). Data are means ± SEM; \*\**P* < 0.01; n.s, non-significant.

### CCK Protects Human Pancreatic β-cells from Apoptosis *In Vitro*

Human islets express CCK and both CCK receptors, although at highly variable levels (Fig.3A) ^30,33^ suggesting that they may also be amenable to protection from CCK receptor-mediated treatments. Intact human islets from deceased organ donors were pretreated with 100 nM CCK-8 or vehicle for 24 hours and then treated with pro-inflammatory human cytokine cocktail. Human islet donor information in Appendix Table 1. β-cells from dispersed human islets treated with CCK-8 before cytokine cocktail exposure had significantly reduced TUNEL staining in comparison to those treated with vehicle before cytokines (1.81-fold ± 0.21, *p<0.05)* (Fig.3B). CCK-8 treatment also decreased apoptosis in dispersed human islet cells, as measured by cytokine-induced caspase 3/7 activity (Fig. 3C). Together, these results demonstrate the efficacy of CCK treatment in promoting β-cell survival in human islets.

**Fig 3.**
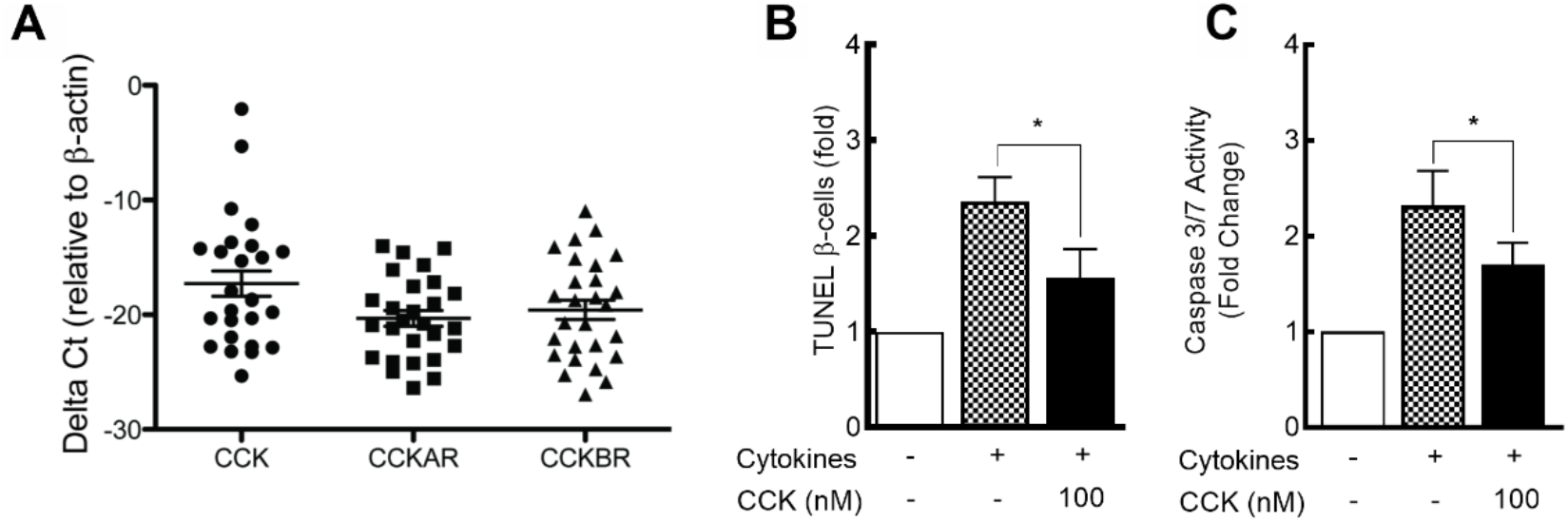
CCK protects human islets from cytokine-induced apoptosis. Human islets express CCK, CCKAR, and CCKBR mRNA at variable levels (*n* = 25) (A). Fold increase of TUNEL-positive/insulin-positive cells (B), and caspase 3/7 activity (C) of human islets treated with cytokine cocktail is reduced when co-treated with CCK. Data are means ± SEM (*n* = 5); \**P* < 0.05.

### Systemic CCK-8 Does Not Alter Body Weight or Blood Glucose Levels in Mouse

To provide evidence that CCK treatment could protect human islets *in vivo*, we transplanted a sub-therapeutic number of human islets (Appendix Table 1) under the kidney capsule of immunodeficient (NOD/SCID) mice with streptozotocin (STZ)-induced diabetes and treated these mice with CCK or saline via an osmotic pump for 3 weeks. Human islets were pre-treated with CCK or vehicle control for 24 hours before transplant. In our xenograft model we transplanted less than 1,000 islet equivalents (IEQs) of human islets per mouse, which we predicted would be insufficient to restore euglycemia. Our goal was to keep the islets in a metabolically unfavorable environment of persistent hyperglycemia, similar to that seen in human diabetes. The amount of CCK delivered by osmotic pump was calculated to approximate the daily dose previously administered to obese mice by twice daily injection^28^. We confirmed that the animals that received the CCK treatment had increased levels of circulating serum CCK and achieved absolute serum concentrations of approximately 20 pM, which is only modestly higher than the physiologic postprandial levels of CCK in humans^41,42^ (Fig.4A). However, blood glucose levels (Fig.4B) and body weight (Fig.4D) of the human islet transplant recipient animals receiving CCK-8 infusion were unaltered by circulating CCK. Finally, circulating human insulin was measured and was not different in the two treatment groups (Fig.4C). Since systemic circulation of CCK-8 did not alter the body weight or blood glucose levels in comparison to the control group, any changes in the viability or apoptotic events of transplanted human islets would not have been the result of differences in insulin demand or glucotoxicity among different transplant groups, but most likely a direct effect of CCK-8 on the human islet cells.

**Fig 4.**
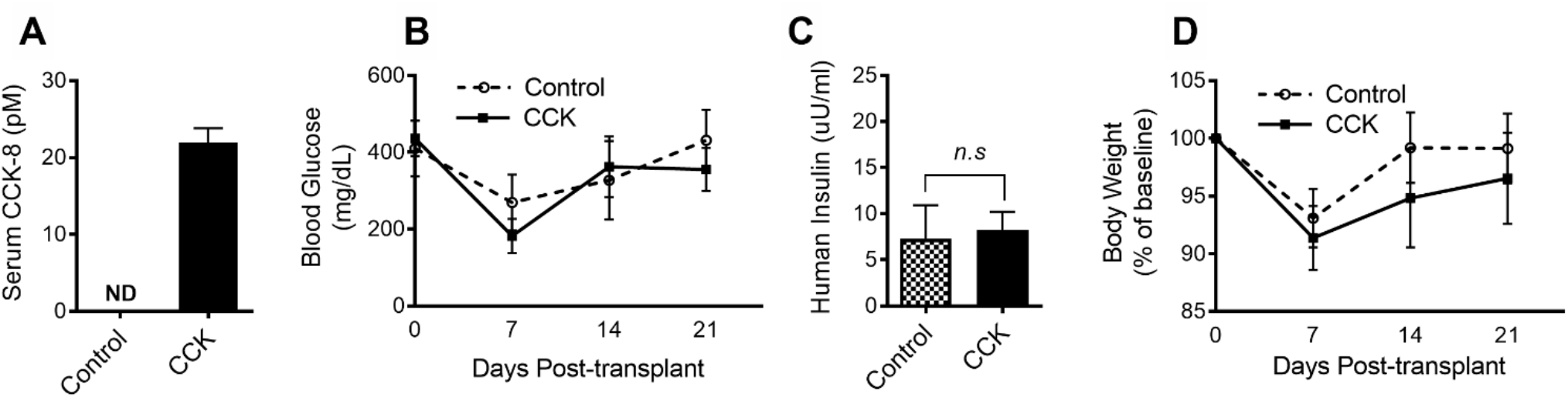
Treatment with CCK does not alter body weight or blood glucose of diabetic mice. Circulating levels of CCK (A) are increased in CCK-treated mice 21 days after transplant. Non-fasted blood glucose levels (B) measured weekly during 21 days post-transplant do not differ between treatment groups. Human insulin (C) is detected in the serum of mice 21 days after transplant and does not differ between control and CCK-treated mice. Body weight was measured weekly and used to calculate percent change (D) during 21 days post-transplant, no significant differences were detected. Data are means ± SEM (*n* = 4-6). ND, not detectable; n.s, non-significant.

### CCK Protects Human Pancreatic β-Cells from Apoptosis Following Transplant

We measured β-cell apoptosis in the islet grafts three-weeks after transplantation. We found reduced TUNEL-positive β-cells in transplanted human islet grafts from mice with CCK treatment (0.5%) in comparison to islet grafts from saline-treated mice (1.61%). This suggests that exogenous CCK treatment protects human β-cells from apoptosis (Fig.5A&B). Taken together, our study indicates that *in vivo*, CCK protects human β-cells from apoptosis under diabetic conditions and in the transplant setting.

**Fig 5.**
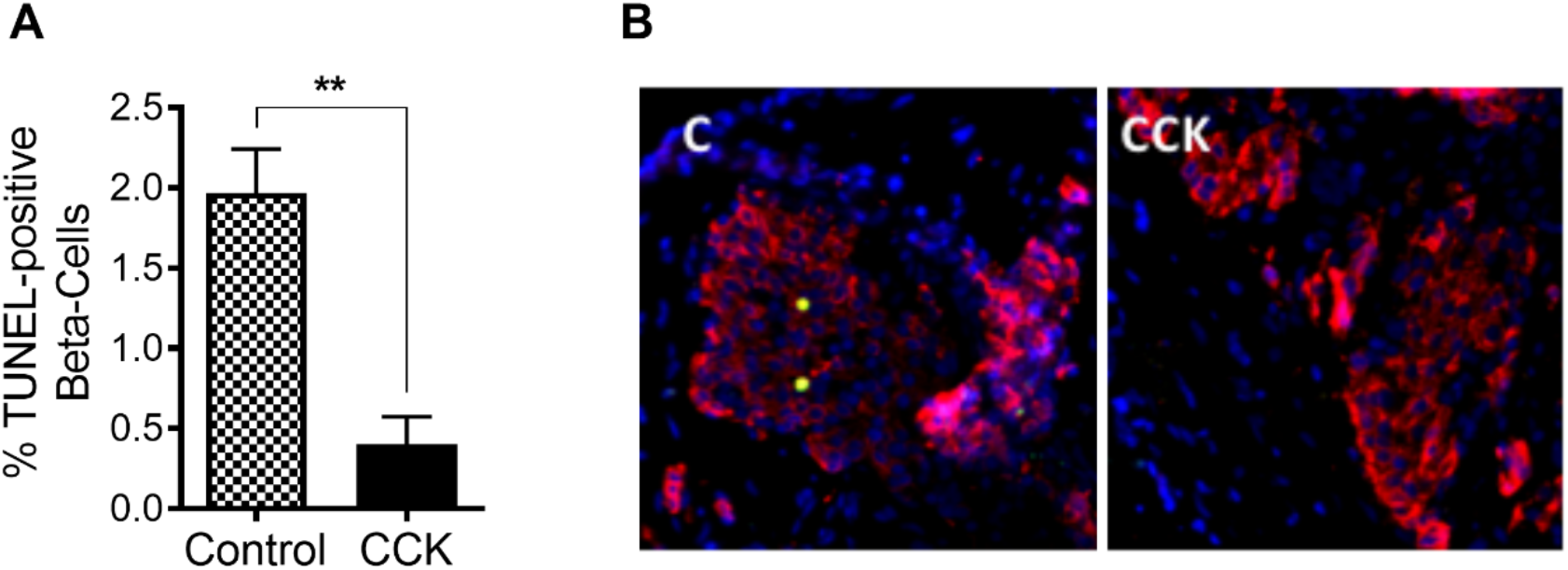
Human islet grafts in mice receiving CCK treatment contain fewer apoptotic β-cells. Quantitative analysis (A) reveals a reduced percentage of TUNEL-positive insulin-positive cells in human islet grafts from CCK-treated mice 21 days after transplant. Representative images of islet grafts (B) from control (left) and CCK-treated (right) transplant mice. DAPI (blue), insulin (red), and TUNEL (green). Data are means ± SEM (*n* = 3-5); \*\**P* < 0.01.

## DISCUSSION

Pancreatic islet transplantation represents a potential cure for diabetes. The fulfillment of this goal is ultimately contingent on achieving long-term survival of the islet graft. Allogenic-ICT can be a curative therapy for T1D, while auto-ICT can preserve the endocrine function of patients undergoing pancreatectomy. However, the major problem with any islet transplantation, particularly for T1D, is a significant loss of islet viability both early and late after the transplant ^**12,43–45**^. In T1D patients, only ~50% of successful islet transplant recipients maintain insulin independence after two years. The loss of islets in the early stages after a transplant requires a high number of islets to restore glucose homeostasis in the patient, often requiring more than one donor per recipient ^**46**^. Inflammation and autoimmunity are one of the causes for the initiation of β-cell destruction during the development of T1D. Transplanted pancreatic islets themselves can also release pro-inflammatory cytokines, which play a significant role in cell failure and death, during and post-islet transplantation ^**43,44,46**^. Thus, identifying factors that can help improve the preservation of pancreatic β-cell mass is in demand to improve islet transplant outcomes.

In rodents, administration of the stable CCK analog (pGlu-Gln-CCK-8) protects mice from obesity-induced diabetes in both high fat-fed and leptin-deficient models ^27,28^. Notably, a study by Irwin et al. ^28^ shows that twice-daily injection of pGlu-Gln-CCK-8 (25 nmol/kg body weight) in high fat-fed mice and *ob*/*ob* mice reduced body weight, improved glucose tolerance, improved insulin sensitivity, and lowered non-fasting glucose, demonstrating the potential of a CCK receptor agonist as an anti-obesity/anti-diabetic agent. However, this study has inherent limitations in isolating the direct beneficial effect of CCK receptor agonism in β-cell survival. Since CCK treated animals had improved glucose tolerance and insulin sensitivity this can indirectly reduce the diabetogenic stress in the islets. Additionally, the high fat diet feeding in their model did not induce the typical compensatory increase in β-cell mass. While these authors reported increased β-cell turnover with CCK treatment, a direct effect of CCK on β-cell survival could not be assessed in this model. In our current study, we demonstrate direct *ex vivo* effects of CCK to reduce β-cell apoptosis and also show reduced apoptosis of human β-cells *in vivo* despite persistent hyperglycemia that is comparable to controls.

This current study demonstrates the relevance of islet CCK signaling in humans by showing that human pancreatic islets express CCK, CCKAR, and CCKBR (Fig.3A), albeit at highly variable expression levels. This is distinct from mouse islets, which primarily express CCKAR with very low CCKBR expression ^**31–34**^. Therefore, this difference in CCK receptor expression is important in assessing species-specific effects of CCK-based therapies on the islet. CCK-8 peptide can activate both CCKAR and CCKBR. We demonstrate that the CCK-8 analog directly protects human islets from cytokine-induced apoptosis (Fig.3B&C). While these *in vitro* studies were encouraging, it was essential to study this effect under *in vivo* settings. We find that continued CCK-8 infusion for three weeks successfully reduced human β-cell apoptosis by over threefold in comparison to the vehicle treated group following the transplant (Fig.5). Not only does this support the promise of CCK-based therapy to promote β-cell survival in human islets, but it also demonstrates that CCK is effective in protecting from the pathophysiologic stressors relevant in diabetes and transplantation such as hyperglycemia, hypoxia, and ER and oxidative stress.

It is also important to note that in this study, we show that CCK-mediated protection is occurring in a pharmacologically validated mechanism via receptor-mediated effects. We achieve this by first demonstrating a dose-response relationship of the protective effects of CCK in INS1E cells across the pM to uM range (Fig.1B). The wide range of effective concentrations provides a hint that CCK treatment to target β-cell survival has a large therapeutic index. Using CCK at lower doses will likely minimize the gastrointestinal side effects that impeded progress in prior clinical trials. Notably, the protection of human islets *in vivo* was achieved with circulating levels of CCK only in the 20 pM range. Additionally, we show that this protection disappears in islets from mice devoid of both CCK receptors (Fig.2C), confirming a direct, receptor-mediated effect.

The results we present from the transplant study are encouraging, but there are some intrinsic limitations. We acknowledge that, due to the design of the study including pre-treatment of islets with drug, having one drug dose, and only one late endpoint for islet graft harvesting, it is impossible to determine whether the protection from apoptosis we see in our transplant model is due to the CCK treatment of the islets before the transplant, the systemic CCK infusion, or a combination of the two. We also do not know if the protective effects of CCK treatment are most pronounced in the early post-transplant period or later in response to persistent hyperglycemic stress. To fully harness the protective effects of CCK and reduce unwanted side effects, it will be essential to determine the appropriate timing and dose of CCK treatment necessary to provide adequate β-cell protection.

All experiments in the current study were done using a non-biased CCK analogue that presumably results in pharmacological activation of both CCK receptors. CCK signals through the G-protein coupled receptors, the CCK-A receptor (CCKAR or CCK1R) or CCK-B receptor (CCKBR or CCK2R), to activate many different signal transduction pathways ^**36,47–49**^. Previously, others have demonstrated the direct effect of CCK on the survival of Min6 cells against apoptosis in response to hyperglycemia and suggested that this effect depended on β-arrestin-mediated signaling ^**36**^. However, the CCK receptor signaling pathways employed by β-cells are not fully understood. Since human islet cells express both the CCKA receptor and the CCKB receptors, future studies will be needed to address specific CCK signaling pathways in the human endocrine pancreas. Identifying the specific CCK receptor subtype activation that affords protection against β-cell apoptosis in human islets in our transplant model will allow further development of CCKR agonists to improve transplant outcomes with minimal adverse effects.

In conclusion, we demonstrate here, for the first time, that CCK protects human β-cells from apoptosis. We provide evidence through *in vitro*, *ex vivo*, and *in vivo* studies using isolated human islets to suggest that this protection occurs through direct effects on the β-cell and in response to several forms of apoptotic stress. CCK has been studied as a potential therapeutic measure against obesity, due to its CNS directed effects on satiety. Our work demonstrates that CCK also has direct positive effects on protecting human β-cell mass, adding to its benefits for utilization in diabetes therapy. The endogenous function of CCK in human pancreatic islet remains an open question. The role of CCK in intra-islet autocrine and paracrine signaling are suggested ^**32**^and other studies also show its potential role in exocrine-endocrine crosstalk for islet mass expansion and regulation in the context of diabetic mice models ^**35**^. Future studies will be essential to discover the CCK signaling pathways activated in β-cells and explore how these pathways can be targeted to develop an effective therapy with few adverse effects.

## MATERIALS AND METHODS

### Viability Assay

INS1E rat insulinoma cells were cultured in RMPI 1640 (Thermo Fisher Scientific, #11875093) with 2.05 mM Glutamax (Cellgro, 35050079) supplemented with 5% (v/v) heat-inactivated FBS, 1% P/S, 10 mM HEPES, 1 mM sodium pyruvate, and 50 μM freshly added beta-mercaptoethanol (Sigma, M7522) at 37 °C and 5% CO2. INS1E cells were seeded in 12 well plates at a density of 0.1 × 10^6^ cells/well and were incubated for 24 hours before any treatment. Cells were then incubated in fresh media containing either 100 nM CCK or 100 nM saline vehicle control up to 72 hours with concurrent treatment with a mouse cytokine cocktail containing 50-ng/mL TNFα (Miltenyi Biotec, #139-101-687), 10-ng/mL IL-1β (Miltenyi Biotec, #130-101-680), and 50 ng/mL IFN-γ (Miltenyi Biotec, #130-105-785) as described ^50^. Cell media containing the treatment was refreshed at 36 hours of incubation for the time course study groups that were treated for more than 24 hours. At each time point, the growth media containing the suspended cells was collected and centrifuged and combined with the adherent cells that were released with 0.25% trypsin (Sigma, 59428C) and then resuspended in 100 ul of growth media containing FBS. 10 ul of suspended cells were then added to a 10ul solution of 0.4% trypan blue in a buffered isotonic salt solution (Bio-Rad, #1450021), pH 7.3, and measured for viability using TC10 automated cell counter (Bio-Rad, #145-0010).

### Apoptosis Assay – Imaging Flowcytometry

Apoptosis was measured in INS1E cells and islets from wild-type (WT) and CCK receptor KO mice treated with cytokine cocktail and CCK for 24 hours. Cultured islets or cells were transferred to a 15 mL conical tube with the incubated media. The culture plate was rinsed with 2 mL PBS and add this to the tube to ensure complete transfer. Islets and cells were pelleted at *800xg* for 3 min at 4°C followed by 1 minute wash using cold PBS. Islets and cells were resuspended in 1 ml Sigma Dissociation Solution (Sigma), then shaken horizontally in a 37°C water bath for 8 min at 100 RPM followed by placement of the tube on ice and addition of 2 mL of media to stop the dissociation process. Cells were then gently disrupted and incubated on ice for another 5 minutes before pelleted again at low speed for 5 min at 4°C. Supernatant was removed and the pellet was resuspended in 1x Annexin Binding Buffer containing 0.5% BSA. 5x Annexin binding buffer was made with 50 mM HEPES, 700 mM NaCl, 12.5 mM CaCl2, in water filled to 50 ml. The cells were then pipetted up and down for 30x, then 1 μl of this cell suspension was placed on a microscope slide to check for dissociation. Depending on the level of clumping, the cells were pipetted up and down for another 20-30x more times or until the majority of the cells were single cells. Cells were then counted on a hemocytometer and were then resuspend in 1x Annexin Binding Buffer containing 0.5% BSA to a final concentration of 1×10^6^ cells/ml. An aliquot of 100 μl of cells (1×10^5^ cells) was mixed with Propidium Iodide (50 μg/mL) and Annexin V-AF488 (Invitrogen #A13201). Unstained and individually stained control samples were prepared in separate tubes. All tubes were incubated at room temp in the dark for 15 min. Samples were diluted immediately before reading with the addition of 400 ul 1x Annexin Binding Buffer containing 0.5% BSA. Imaging Flow data was acquired by imaging flow cytometer (Amnis EMD Millipore, Image Stream Mark II). During the analysis, 10,000 live cell images were captured at 40x. Data were analyzed using ImageStream (Image Stream®X, Amaris) for the quantification of propidium iodide and annexin V stains that indicate the different stages of apoptosis.

### Ex vivo Dispersed Islet TUNEL and Caspase 3/7 Activity Assays

Using serum-free media supplemented with 5 g/l BSA fraction V (Roche, #107351080001), islets were pre-treated with 100 nM CCK, saline vehicle control for 24 hours, before additional 24 hours of cytokine or 10 uM thapsigargin treatment to induce apoptosis. Human islets were treated with a cytokine cocktail containing human 1,000 U/ml TNFα, 75 U/ml IL-1β, and 750 U/ml IFN-γ (PeproTech, #200-01B, #300-02, #300-01A). After treatments, islets were dispersed, plated on Poly-L-lysine pre-coated glass coverslips and fixed. β-cell apoptosis was measured using TUNEL (Promega, #G3250) followed by insulin staining (Dako, A0564). Imaging was performed using an EVOS FL Autofluorescence microscope (Life Technologies). ImageJ (NIH) or Adobe Photoshop (Adobe) was used to count DAPI (Vector Labs, H-1200-10) in at least 9 randomly chosen fields per treatment group for each replicate. Dispersed human islets were plated in a 96 well plate at 20,000 cells per well and incubated for 24 hours prior to treatment. For Figure 3C, dispersed islets were pre-treated with 100 nM CCK or vehicle control for 1-hour prior to treatment with human cytokines as above. Caspase 3/7 activity was measured 24 hours post-cytokine treatment using the Caspase-Glo® 3/7 Assay System (Promega, #G8090).

### Animals, Islet Isolation and Culture

Animal care and experimental procedures were performed with approval from the University of Wisconsin Animal Care and Use Committee to meet acceptable standards of humane animal care. Male C57BL/6J (~10 to 15-week-old) (JAX Stock Number #000664) and NOD/SCID (9-week-old) mice (JAX Stock Number #001303) were purchased from The Jackson Laboratory for islet isolation and transplant studies, respectively. CCK receptor double knockout mice, 129-*Cckar*^tm1Kpn^ *Cckbr*^tm1Kpn/J^ (JAX Stock Number #006365) ^51^ age 14-20 weeks were used in the studies. These mice are in the 129S1/SvImJ background (JAX Stock Number #002448) and were obtained from Jackson Laboratories after cryorecovery. Mice were housed in facilities with a standard light-dark cycle and fed ad libitum. Mouse pancreatic islets were isolated using collagenase digestion and hand-picked as previously described ^52,53^. Isolated islets were cultured at 37°C and 5% CO_2_ in RPMI 1640 media (Thermo Fisher Scientific, #11879020) containing 8 mM glucose, supplemented with 10% heat-inactivated fetal bovine serum (FBS), 100 units/ml penicillin and 100 μg/ml streptomycin (1% P/S) (Thermo Fisher Scientific).

### Human islet culture and mRNA Levels

Human islets were obtained through the Integrated Islet Distribution Program (IIDP). Upon arrival, islets were handpicked and then cultured in RPMI 1640 media as described above but without antibiotics and antimycotics. Islets were cultured overnight to confirm viability and sterility before treatments. Islets were cultured up to 7 days, and the media was renewed every other day. After culture, RNA was isolated, and gene expression was quantified via quantitative PCR with SYBR and primers for human *CCK (hsCCK forward* TGA GGG TAT CGC AGA GAA CGG ATG, hsCCK *reverse* TGT AGT CCC GGT CAC TTA TCC TGT), *CCKAR (hsCCKAR forward* TGG AAG CAA CAT CAC TCC TC, hsCCKAR *reverse* CAC GCT GAG CAG GAA TAT CA), and *CCKBR (hsCCKBR forward* GAT GTG GTT GAC AGC CTT CT, hsCCKBR *reverse* GGG CTG ATC CAA GCA GAA A) normalized to Beta-Actin (*forward* TCA AGA TCA TTG CTC CTG AGC, *reverse* TCA AGA TCA TTG CTC CTG AGC).

### Human Islet Transplantation and Graft Harvest

A modified version of human islet transplantation described by Montanya *et al*. was used for the study ^54^. A single high dose (200 mg/Kg) of streptozotocin (Sigma, #S0130) was administered via intraperitoneal injection two days before transplant to induce hyperglycemia. Hyperglycemia (glucose >300 mg/dL) was confirmed with tail vein blood sample using a glucometer prior to proceeding with transplant. Human islets were cultured in media supplemented with 100 nM CCK (sulfated-(pGlu-Gln)-CCK-8, American Peptide Company) or saline vehicle control for 24 hours before transplant. Confirmed hyperglycemic mice were anesthetized using isoflurane, and approximately 1,000 islet equivalents were placed under the kidney capsule. While our initial study design was to have a paired sample of both CCK-treated and saline-treated from each human islet donor, in some cases we had death of a mouse during or post-surgery or technical difficulties during the transplant that led to insufficient islet delivery in one animal. Therefore, the final data do not always represent paired treatment groups. An infusion pump (Azlet 1004) containing saline or CCK was placed subcutaneously in the back of the mice at the same time as the islet transplant to have continuous CCK infusion (52 pmol/hr) for 3 weeks. We calculated the concentration of infused CCK to recapitulate the theoretical amount of CCK delivered through twice-daily intraperitoneal injection dosing done in studies by Irwin et al. that led to beneficial effects on glucose and weight in obese mice ^27,28^. Mice were monitored during recovery and checked for surgical complications in the post-operative period. Bodyweight and randomly fed blood glucose were measured every 2-3 days following the transplant for a total of three weeks. Blood glucose was measured using a tail nick blood sample and glucometer (Bayer Contour Next EZ). Three weeks post-transplant mice were anesthetized using Avertin (2,2,2-TRIBROMOETHANOL, 97% T48402, 500mg/kg) and terminal serum was collected by cardiac puncture and stored for further assays. Kidneys containing islet grafts were harvested and fixed in 10% formalin (Fisher SF100) for 48 hours. 10% Formalin-fixed kidneys were paraffin-embedded and sectioned for immunofluorescence staining.

### CCK and Human Insulin Measurement

Serum CCK levels were measured three weeks post-transplant using a CCK radioimmunoassay (Alpco Diagnostics - now discontinued) as described by Rehfeld. ^55^. Serum samples were collected through cardiac puncture and plasma extracted by mixing with 96% ethanol and evaporating to dryness using a vacuum centrifuge. The dry extracts were dissolved in Alpco diluent (reagent D) and stored at −20 °C until assayed. The procedure provided by Alpco was followed for the radioimmunoassay. Briefly, samples were mixed with anti-CCK-8 (reagent A) and incubated for 2 days at 2-8 °C followed by the addition of I-CCK-8 (reagent B) and another 4-day incubation at 2-8 °C. Finally, the double antibody solid phase (reagent C) was added, incubated 60 minutes at 2-8 °C, centrifuged, and the supernatant discarded. A gamma counter was used to measure radioactivity with a counting time of 2-4 minutes. Insulin secreted from the islet grafts was measured in the mouse serum three weeks post-transplant using a human insulin ELISA (Millipore, EZHI-14K).

### Immunofluorescence Staining

Paraffin-embedded islet grafts within the kidneys or intact mouse or human islets were stained for insulin using polyclonal guinea pig anti-insulin antibody (Dako, A0564), and apoptosis was measured using the DeadEnd Fluorometric terminal deoxynucleotidyl transferase-mediated deoxyuridine triphosphate nick-end labeling (TUNEL) system (Promega, #G3250). Images from each islet graft were obtained using an EVOS FL Autofluorescence microscope (Life Technologies). ImageJ software was used to quantify total insulin-positive cells and TUNEL-insulin co-staining cells.

### Statistics

Assessment of statistical significance between groups was determined using GraphPad Prism by 2-tailed Student’s t or ANOVA tests as the non-parametric equivalent. Bonferroni posthoc test was performed to correct for multiple comparisons where appropriate. A probability of error less than 5% was considered significant (i.e., P < 0.05). Statistical information for experiments (data representation, P values, and n numbers) can be found in the figure legends. In all panels, data are represented as mean ± SEM.

## ACKNOWLEDGEMENTS

We appreciate assistance from Dr. Sara Sackett from Dr. Jon Odorico’s laboratory (UW-Madison) for training on islet transplant procedures. We thank Dr. Matt Flowers, Grace H. Yang, Sarah E. Nustad for manuscript feedback and Soyoun Kim for appendix data table organization.

## APPENDIX

**Suppl. Table 1.**
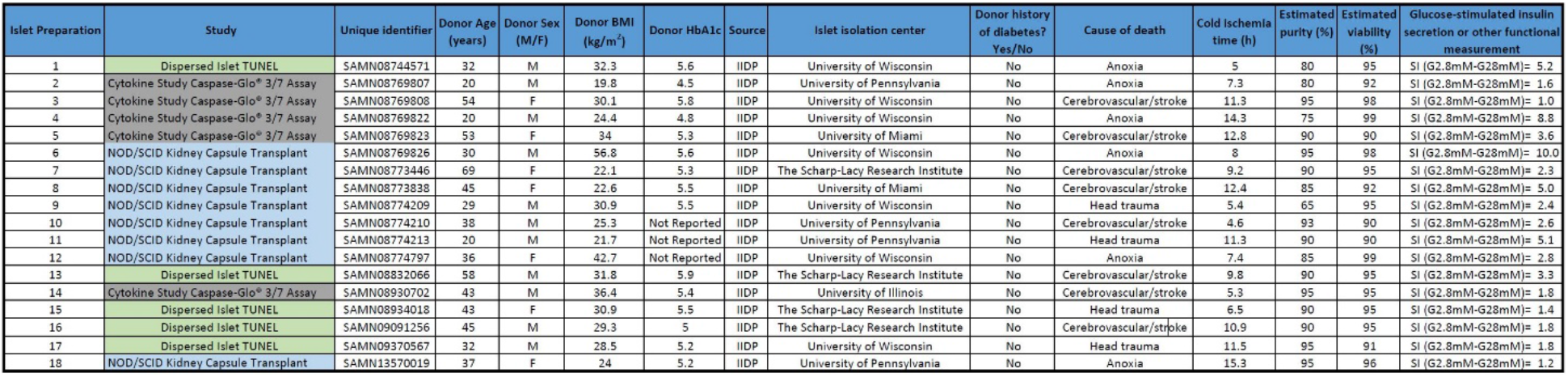
Human islet donors. Adapted from Hart NJ, Powers AC (2018) Progress, challenges, and suggestions for using human islets to understand islet biology and human diabetes. Diabetologia.

## BIBLIOGRAPHY

1. Klöppel G, Löhr M, Habich K, Oberholzer M, Heitz PU. Islet Pathology and the Pathogenesis of Type 1 and Type 2 Diabetes mellitus Revisited. Pathol Immunopath R. 2008;4(2):110–125. doi:10.1159/000156969

2. Butler AE, Janson J, Soeller WC, Butler PC. Increased β-Cell Apoptosis Prevents Adaptive Increase in β-Cell Mass in Mouse Model of Type 2 Diabetes Evidence for Role of Islet Amyloid Formation Rather Than Direct Action of Amyloid. Diabetes. 2003;52(9):2304–2314. doi:10.2337/diabetes.52.9.2304

3. Prentki M, Nolan CJ. Islet β cell failure in type 2 diabetes. The Journal of clinical investigation. 2006;116.

4. Tao Z, Shi A, Zhao J. Epidemiological Perspectives of Diabetes. Cell Biochem Biophys. 2015;73(1):181–185. doi:10.1007/s12013-015-0598-4

5. National Diabetes Statistics Report, 2020. Centers for Disease Control and Prevention, U.S. Dept of Health and Human Services; 2020.

6. Vetere A, Choudhary A, Burns SM, Wagner BK. Targeting the pancreatic β-cell to treat diabetes. Nat Rev Drug Discov. 2014;13(4):278–289. doi:10.1038/nrd4231

7. Sneddon JB, Tang Q, Stock P, et al. Stem Cell Therapies for Treating Diabetes: Progress and Remaining Challenges. Cell Stem Cell. 2018;22(6):810–823. doi:10.1016/j.stem.2018.05.016

8. Song I, Muller C, Louw J, Bouwens L. Regulating the Beta Cell Mass as a Strategy for Type-2 Diabetes Treatment. Curr Drug Targets. 2015;16(5):516–524. doi:10.2174/1389450116666150204113928

9. Narayanan S, Loganathan G, Dhanasekaran M, et al. Intra-islet endothelial cell and β-cell crosstalk: Implication for islet cell transplantation. World J Transplant. 2017;7(2):117–128. doi:10.5500/wjt.v7.i2.117

10. Sutherland DER, Matas AJ, Najarian JS. Pancreatic Islet Cell Transplantation. Surg Clin N Am. 1978;58(2):365–382. doi:10.1016/s0039-6109(16)41489-1

11. Langer RM. Islet Transplantation: Lessons Learned Since the Edmonton Breakthrough. Transplant P. 2010;42(5):1421–1424. doi:10.1016/j.transproceed.2010.04.021

12. Shapiro AMJ, Pokrywczynska M, Ricordi C. Clinical pancreatic islet transplantation. Nat Rev Endocrinol. 2016;13(5):268–277. doi:10.1038/nrendo.2016.178

13. Miyasaka K, Funakoshi A. Cholecystokinin and cholecystokinin receptors. J Gastroenterol. 2003;38(1):1–13. doi:10.1007/s005350300000

14. Zhao X-Y, Ling Y-L, Li Y-G, Meng A-H, Xing H-Y. Cholecystokinin octapeptide improves cardiac function by activating cholecystokinin octapeptide receptor in endotoxic shock rats. World J Gastroentero. 2005;11(22):3405–3410. doi:10.3748/wjg.v11.i22.3405

15. Miyamoto S, Shikata K, Miyasaka K, et al. Cholecystokinin Plays a Novel Protective Role in Diabetic Kidney Through Anti-inflammatory Actions on Macrophage Anti-inflammatory Effect of Cholecystokinin. Diabetes. 2012;61(4):897–907. doi:10.2337/db11-0402

16. Zuelli FM das GC, Cárnio EC, Saia RS. Cholecystokinin protects rats against sepsis induced by Staphylococcus aureus. Med Microbiol Immun. 2014;203(3):165–176. doi:10.1007/s00430-014-0328-3

17. Sui Y, Vermeulen R, Hökfelt T, Horne MK, Stanić D. Female mice lacking cholecystokinin 1 receptors have compromised neurogenesis, and fewer dopaminergic cells in the olfactory bulb. Front Cell Neurosci. 2013;7:13. doi:10.3389/fncel.2013.00013

18. Reisi P, Ghaedamini AR, Golbidi M, Shabrang M, Arabpoor Z, Rashidi B. Effect of cholecystokinin on learning and memory, neuronal proliferation and apoptosis in the rat hippocampus. Adv Biomed Res. 2015;4(1):227. doi:10.4103/2277-9175.166650

19. Nishimura S, Bilgüvar K, Ishigame K, Sestan N, Günel M, Louvi A. Functional Synergy between Cholecystokinin Receptors CCKAR and CCKBR in Mammalian Brain Development. Plos One. 2015;10(4):e0124295. doi:10.1371/journal.pone.0124295

20. Yang S, Ling Y, Ling Y, Duan G, Wang J. Effect of cholecystokinin-octapeptide on focal cerebral ischemia/reperfusion injury in rats. Chinese Journal of Pathophysiology. Published online 1986.

21. Liu Y, Zhang Y, Gu Z, et al. Cholecystokinin octapeptide antagonizes apoptosis in human retinal pigment epithelial cells. Neural Regen Res. 2014;9(14):1402–1408. doi:10.4103/1673-5374.137596

22. Matson CA, Ritter RC. Long-term CCK-leptin synergy suggests a role for CCK in the regulation of body weight. Am J Physiology-regulatory Integr Comp Physiology. 1999;276(4):R1038–R1045. doi:10.1152/ajpregu.1999.276.4.r1038

23. Matson CA, Wiater MF, Kuijper JL, Weigle DS. Synergy Between Leptin and Cholecystokinin (CCK) to Control Daily Caloric Intake. Peptides. 1997;18(8):1275–1278. doi:10.1016/s0196-9781(97)00138-1

24. Matson CA, Reid DF, Ritter RC. Daily CCK injection enhances reduction of body weight by chronic intracerebroventricular leptin infusion. Am J Physiology-regulatory Integr Comp Physiology. 2002;282(5):R1368–R1373. doi:10.1152/ajpregu.00080.2001

25. Lo C, King A, Samuelson LC, et al. Cholecystokinin Knockout Mice Are Resistant to High-Fat Diet-Induced Obesity. Gastroenterology. 2010;138(5):1997–2005. doi:10.1053/j.gastro.2010.01.044

26. Cano V, Merino B, Ezquerra L, Somoza B, Ruiz-Gayo M. A cholecystokinin-1 receptor agonist (CCK-8) mediates increased permeability of brain barriers to leptin. Brit J Pharmacol. 2008;154(5):1009–1015. doi:10.1038/bjp.2008.149

27. Irwin N, Frizelle P, Montgomery IA, Moffett RC, O’Harte FPM, Flatt PR. Beneficial effects of the novel cholecystokinin agonist (pGlu-Gln)-CCK-8 in mouse models of obesity/diabetes. Diabetologia. 2012;55(10):2747–2758. doi:10.1007/s00125-012-2654-6

28. Irwin N, Montgomery IA, Moffett RC, Flatt PR. Chemical cholecystokinin receptor activation protects against obesity-diabetes in high fat fed mice and has sustainable beneficial effects in genetic ob/ob mice. Biochem Pharmacol. 2013;85(1):81–91. doi:10.1016/j.bcp.2012.10.008

29. Irwin N, Pathak V, Flatt PR. A Novel CCK-8/GLP-1 Hybrid Peptide Exhibiting Prominent Insulinotropic, Glucose-Lowering, and Satiety Actions With Significant Therapeutic Potential in High-Fat–Fed Mice. Diabetes. 2015;64(8):2996–3009. doi:10.2337/db15-0220

30. Souza AH de, Tang J, Yadev AK, et al. Intra-islet GLP-1, but not CCK, is necessary for β-cell function in mouse and human islets. Sci Rep-uk. 2020;10(1):2823. doi:10.1038/s41598-020-59799-2

31. Lavine JA, Raess PW, Stapleton DS, et al. Cholecystokinin is up-regulated in obese mouse islets and expands beta-cell mass by increasing beta-cell survival. Endocrinology. 2010;151(8):3577–3588. doi:10.1210/en.2010-0233

32. Lavine JA, Kibbe CR, Baan M, et al. Cholecystokinin expression in the β-cell leads to increased β-cell area in aged mice and protects from streptozotocin-induced diabetes and apoptosis. Am J Physiol-endoc M. 2015;309(10):E819–E828. doi:10.1152/ajpendo.00159.2015

33. Linnemann AK, Neuman JC, Battiola TJ, Wisinski JA, Kimple ME, Davis DB. Glucagon-Like Peptide-1 Regulates Cholecystokinin Production in β-Cells to Protect From Apoptosis. Mol Endocrinol. 2015;29(7):978–987. doi:10.1210/me.2015-1030

34. Lavine JA, Attie AD. Gastrointestinal hormones and the regulation of β-cell mass. Ann Ny Acad Sci. 2010;1212(1):41–58. doi:10.1111/j.1749-6632.2010.05802.x

35. Egozi A, Halpern KB, Farack L, Rotem H, Itzkovitz S. Zonation of Pancreatic Acinar Cells in Diabetic Mice. Cell Reports. 2020;32(7):108043. doi:10.1016/j.celrep.2020.108043

36. Ning S, Zheng W, Su J, et al. Different downstream signalling of CCK1 receptors regulates distinct functions of CCK in pancreatic beta cells. Brit J Pharmacol. 2015;172(21):5050–5067. doi:10.1111/bph.13271

37. Khan D, Vasu S, Moffett RC, Irwin N, Flatt PR. Expression of Gastrin Family Peptides in Pancreatic Islets and Their Role in bgr-Cell Function and Survival. Pancreas. 2018;47(2):190–199. doi:10.1097/mpa.0000000000000983

38. Pathak V, Flatt PR, Irwin N. Cholecystokinin (CCK) and related adjunct peptide therapies for the treatment of obesity and type 2 diabetes. Peptides. 2018;100(Diabetes Res. Clin. Pract. 87 3 2010):229–235. doi:10.1016/j.peptides.2017.09.007

39. Engin F. ER stress and development of type 1 diabetes. J Invest Med. 2016;64(1):2. doi:10.1097/jim.0000000000000229

40. Brozzi F, Nardelli TR, Lopes M, et al. Cytokines induce endoplasmic reticulum stress in human, rat and mouse beta cells via different mechanisms. Diabetologia. 2015;58(10):2307–2316. doi:10.1007/s00125-015-3669-6

41. Liddle RA, Morita ET, Conrad CK, Williams JA. Regulation of gastric emptying in humans by cholecystokinin. J Clin Invest. 1986;77(3):992–996. doi:10.1172/jci112401

42. Gibbons C, Finlayson G, Caudwell P, et al. Postprandial profiles of CCK after high fat and high carbohydrate meals and the relationship to satiety in humans. Peptides. 2016;77:3–8. doi:10.1016/j.peptides.2015.09.010

43. Li X, Meng Q, Zhang L. The Fate of Allogeneic Pancreatic Islets following Intraportal Transplantation: Challenges and Solutions. J Immunol Res. 2018;2018:1–13. doi:10.1155/2018/2424586

44. Narayanan S, Bhutiani N, Adamson DT, Jones CM. Pancreatectomy, Islet Cell Transplantation, and Nutrition Considerations. Nutr Clin Pract. Published online 2020. doi:10.1002/ncp.10578

45. Warnock GL, Kneteman NM, Ryan EA, Rabinovitch A, Rajotte RV. Long-term follow-up after transplantation of insulin-producing pancreatic islets into patients with Type 1 (insulin-dependent) diabetes mellitus. Diabetologia. 1992;35(1):89–95. doi:10.1007/bf00400857

46. Lehmann R, Spinas GA, Moritz W, Weber M. Has Time Come for New Goals in Human Islet Transplantation? Am J Transplant. 2008;8(6):1096–1100. doi:10.1111/j.1600-6143.2008.02214.x

47. Hansen T v.O. Cholecystokinin gene transcription: promoter elements, transcription factors and signaling pathways. Peptides. 2001;22(8):1201–1211. doi:10.1016/s0196-9781(01)00443-0

48. Dufresne M, Seva C, Fourmy D. Cholecystokinin and Gastrin Receptors. Physiol Rev. 2006;86(3):805–847. doi:10.1152/physrev.00014.2005

49. Zeng Q, Ou L, Wang W, Guo D-Y. Gastrin, Cholecystokinin, Signaling, and Biological Activities in Cellular Processes. Front Endocrinol. 2020;11:112. doi:10.3389/fendo.2020.00112

50. Graham KL, Fynch S, Pappas EG, Tan C, Kay TW, Thomas HE. Isolation and culture of the islets of Langerhans from mouse pancreas. European Journal of Immunology. Published online 2015.

51. Kopin AS, Mathes WF, McBride EW, et al. The cholecystokinin-A receptor mediates inhibition of food intake yet is not essential for the maintenance of body weight. J Clin Invest. 1999;103(3):383–391. doi:10.1172/jci4901

52. Carter JD, Dula SB, Corbin KL, Wu R, Nunemaker CS. A Practical Guide to Rodent Islet Isolation and Assessment. Biol Proced Online. 2009;11(1):3. doi:10.1007/s12575-009-9021-0

53. Rabaglia ME, Gray-Keller MP, Frey BL, Shortreed MR, Smith LM, Attie AD. α-Ketoisocaproate-induced hypersecretion of insulin by islets from diabetes-susceptible mice. Am J Physiol-endoc M. 2005;289(2):E218–E224. doi:10.1152/ajpendo.00573.2004

54. Estil·les E, Téllez N, Nacher M, Montanya E. A Model for Human Islet Transplantation to Immunodeficient Streptozotocin-Induced Diabetic Mice. Cell Transplant. 2018;27(11):1684–1691. doi:10.1177/0963689718801006

55. Rehfeld JF. Accurate measurement of cholecystokinin in plasma. Clin Chem. 1998;44(5):991–1001. doi:10.1093/clinchem/44.5.991

